# Theta activity discriminates high-level, species-specific body processes

**DOI:** 10.1101/2023.11.13.566536

**Authors:** Jane Chesley, Lars Riecke, Juanzhi Lu, Rufin Vogels, Beatrice de Gelder

## Abstract

Among social stimuli that trigger rapid reactions, body images occupy a prominent place. Given that bodies carry information about other agents’ intentions, actions and emotional expressions, a foundational question concerns the neural basis of body processing. Previous fMRI studies have investigated this but were not yet able to clarify the time course and its functional significance. The present EEG study investigated the role of slow oscillatory cortical activity in body processing and species-specificity. Human participants viewed naturalistic images of human and monkey bodies, faces, and objects, along with mosaic-scrambled versions to control for low-level visual features. Analysis of event-related theta power (4 – 7 Hz) combined with data-driven methods revealed a strong, body-evoked neural response that is specific to human bodies and likely originates from a widespread cortical region during a time window of 150 – 550 ms after the onset of the body image. Our results corroborate recent research proposing a widespread, species-specific cortical network of human body processing. We submit that this network may play an essential role in linking body processes to movement intentions.

## 1. Introduction

Social species vitally rely on information from their conspecifics to navigate the natural and social world. During social interactions, humans rapidly decode cues from the faces and bodies of others, which hold information relevant to identity, emotions, and actions. While the role of faces in regulating social interactions has been well-established (Freiwald et al., 2016; Powell et al., 2018; Schwiedrzik et al., 2015), evidence for a role of whole-body processing is still accumulating. Body-selective areas were first reported in the lateral occipitotemporal cortex (LOTC), termed the extrastriate body area (EBA) and fusiform body area (FBA) (Downing et al., 2001; Peelen & Downing, 2005). Further research has reported body-selective responses widespread throughout the brain in the posterior superior temporal sulcus (pSTS) (Kret et al., 2011; Candidi et al., 2015), temporoparietal junction (TPJ), frontal cortex and parietal motor areas (Pichon et al., 2009), as well as subcortical areas (de Gelder & Poyo Solanas, 2021, Poyo Solanas et al., 2020; Swann et al., 2012). Furthermore, recent research combining advanced data-driven methods with 7-Tesla functional magnetic resonance imaging (fMRI) has revealed two large-scale networks widespread throughout the right STS and lateral occipital cortex (LOC) that are specifically selective for human body stimuli, suggesting that body processing may be species-specific (Li et al., 2023).

Additional lines of research using electroencephalography (EEG) have investigated the millisecond-precise timing of neural responses to bodies. With this method, event-related potential (ERP) studies have reported that, like faces, bodies are processed configurally, as shown by enhanced and delayed body-sensitive N170 ERPs to inverted versus normally oriented bodies (Stekelenburg & de Gelder, 2004). In addition, like faces, emotional information from body stimuli is rapidly encoded in early stages of visual processing, as differences between fearful and neutral body responses can emerge as early as 112 ms after stimulus onset (van Heijnsbergen et al., 2007). A body-specific ERP modulation has consistently been observed at 190 ms post-stimulus (N190) over occipito-temporal regions in response to silhouettes of normal bodies compared to scrambled silhouettes (Thierry et al., 2006) as well as to headless naturalistic bodies compared to plants (Taylor et al., 2010; Moreau et al., 2018), providing further evidence for body-specific processes. Furthermore, intracranial local field potentials (iLFPs) have shown body-selective responses emerging from EBA at 190 ms post-stimulus, with a peak at 260 ms (Pourtois et al., 2007).

While EEG research has consistently shown body-related effects on stimulus-evoked broadband cortical responses, effects on oscillatory cortical responses have been investigated much less. Frequency-specific (narrow-band) oscillatory activity is thought to represent different areal and interareal processing mechanisms (Fries, 2009, 2015; Wang, 2010), and modulations of oscillatory activity have been implicated in various cognitive functions like cognitive control, learning, memory and action regulation (Cavanagh & Frank, 2014; Herweg et al., 2020; Trujillo & Allen, 2007). In particular, neural activity in the theta band (4 – 7 Hz) has been linked to body processes: differential theta activation has been observed over occipito-temporal and pre-frontal regions for body versus face processing within 250 – 500 ms post-stimulus (Bossi et al., 2020). Moreover, these regions have been shown to synchronize their theta activity in the aforementioned time window during the processing of visual body information during social interactions (Moreau et al., 2020). Furthermore, widespread theta activity has been observed throughout the brain within the first 400 ms of stimulus onset for self-and non-self body responses (Çelik et al., 2021). Overall, these findings suggest that oscillatory theta activity within 500 ms after body-image onset might play a relevant role in body processing.

An important methodological challenge in the study of neural representations of bodies is the control of low-level sensory information. Naturally, visual stimuli convey low-and high-level information. Low-level features include elementary visual information of luminance, contrast, and textures, among others (Koch & Ullman, 1987; Veale et al., 2017). On the other hand, high-level features refer to semantic and categorical information, such as the identification of a stimulus as a “body”, “face”, or “object” (Groen et al., 2017; Kandel et al., 2014). An effective approach to isolating the high-level processes in the brain is to include scrambled stimuli in the experimental design, as scrambled stimuli can preserve several low-level stimulus features while destroying higher-level information. Some ERP studies have used scrambled stimuli (van Heijnsbergen et al., 2007), but currently in the field, no oscillatory body research (see above) has adequately controlled for the contributions of low-level visual features with the use of scrambled body stimuli, leaving unclear whether their findings reflect visual or more abstract body representations. The present study aims to bridge this gap by including mosaic-scrambled stimuli that control for low-level features of luminance, contrast, and texture to better understand the role of oscillatory theta activity in high-level body processes.

By using EEG and a data-driven approach, we first identified a strong theta response in a widespread, bi-lateral region within 200 – 550 ms after the onset of visual categorical stimuli. Using an experimental design comprising category conditions (body, face, and object), visual controls (scrambled versions of the categorical stimuli), and species (human and monkey), we then tested whether these responses are human body-specific, while controlling for low-level visual features. Based on previous fMRI research suggesting a large-scale, species-specific cortical network for human body processing (Li et al., 2023), we expected the high-level (scramble-controlled) representations of bodies to be species-specific, with a clear enhancement of human (versus monkey) body processing.

## 2. Methods

### 2.1 Participants

Thirty healthy, right-handed participants with normal or corrected-to-normal vision were recruited for this study. All participants reported no history of psychiatric or neurological disorders. Written consent was obtained from participants prior to the experiment. Participants were compensated in either monetary vouchers or credit points. One participant’s data were excluded from the analysis because she/he presumably misunderstood the attention task (as shown by 0% accuracy); the remaining 29 participants had an average accuracy of 96 ± 4% (mean ± SD) (range = 85 – 100%). Hence, 29 participants’ data were included in the analysis (17 females; age range = 18-37 years; mean age = 23). Procedures were approved by the Ethical Committee of Maastricht University and were in accordance with the Declaration of Helsinki.

### 2.2 Stimuli

Grayscale, naturalistic images of bodies, faces and objects were used as stimuli in the experiment (Fig. 1A). Body and face stimuli were from a human or a monkey. Object stimuli were divided into two sets such that the aspect ratio matched human bodies (set 1) or monkey bodies (set 2). Body stimuli had face information removed with Gaussian blurring. Stimuli were embedded in a white noise background and presented centrally on the computer screen. The size of the stimuli was 9 * 9 degrees of visual angle for human faces, 9 * 20 degrees for human bodies and objects, and 16 * 16 degrees for monkey faces, bodies, and objects.

**Figure 1.**
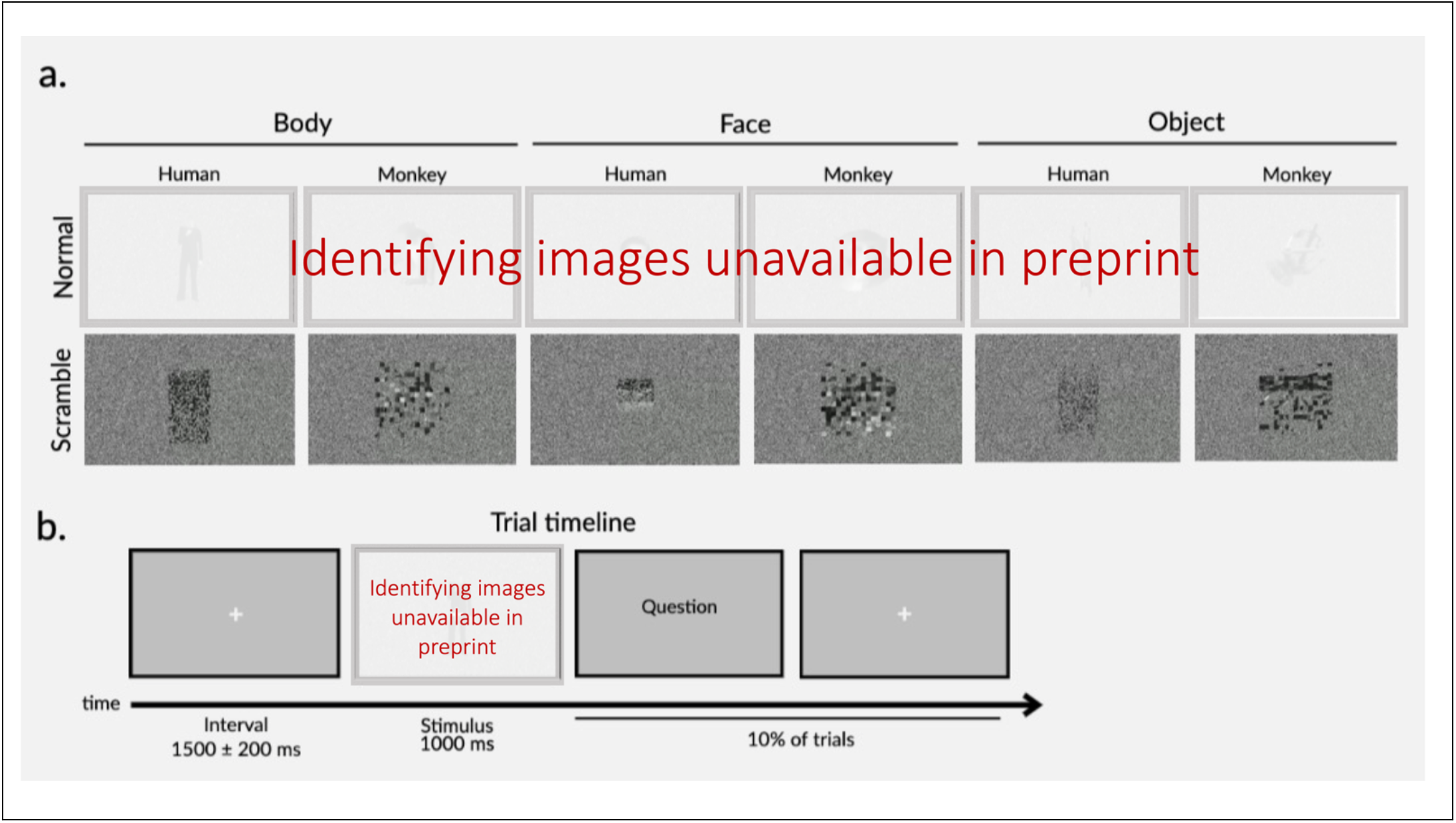
Example stimuli for all conditions (A) and trial timeline (B).

To control for the contribution of low-level visual features, mosaic-scrambled images were included. Mosaic-scrambled images destroyed the whole shape of each body/face/object stimulus, but preserved the low-level features of luminance, contrast, texture, and non-background area (Bognár et al., 2023). This resulted in a total of twelve experimental conditions (human/monkey * body/face/object * normal/scrambled). There were ten different stimuli per condition, which resulted in 120 unique images. All images were adapted from video stimuli used in a previous body perception study (Li et al., 2023; see also Bognár et al., 2023; Kret et al., 2011; Zhu et al., 2013). The images for the present study were extracted from the midpoint (frame 30) of each original video (60 fps). Detailed descriptions of the stimuli can be obtained from the aforementioned papers. Image extraction and stimulus presentation were programmed in MATLAB 2021a (The Mathworks, Natick, MA, USA) with the Psychophysics Toolbox extensions (Brainarrd, 1997; Pelli, 1997; Kleiner et al., 2007) as well as custom code.

### 2.3 Experimental design, task and procedure

The experiment consisted of two experimental sessions, one of which presented images (see Stimuli) and the second of which presented videos of the same stimuli. The order of the two experimental sessions was randomized across participants. The present paper reports the methods, analysis, and results of the former, image-related experimental session; the latter was used for another project.

The main experiment employed a randomized design. There were four runs, all lasting around 6 minutes. During each run, 120 unique images (12 conditions × 10 stimuli; see Stimuli) were presented once in random order. This resulted in a total of four repetitions per stimulus and 40 repetitions per condition. Each trial began with a white fixation cross centered on a gray screen (Fig. 1B). To reduce the temporal expectancy of stimulus presentation, the intertrial interval was jittered at 1500 ms (1500 ± 200 ms). Participants viewed the images on a computer screen (1920 × 1080) at 65 cm from their eyes. A white fixation cross was centered and overlaid on each image. Participants were asked to focus their gaze on the fixation cross and focus their attention on each stimulus. To maintain attention, a question appeared on a random 10% of trials. The question asked about the content of the preceding stimulus (E.g. “What did the previous image show?”), and participants were asked to respond with a button press from a selection of “Body”, “Face”, “Object” or “None of the above.”

### 2.4 EEG acquisition

EEG signals were acquired from 33 electrodes embedded in a fabric cap (EASYCAP GmbH) and arranged in accordance with the international 10-20 system. Scalp electrodes included: AFz, Fz, FCz, Cz, CPz, Pz, Oz, Fp1, Fp2, F3, F4, F7, F8, FC3, FC4, FT7, FT8, C3, C4, T7, T8, CP3, CP4, TP7, TP8, TP9, TP10, P3, P4, P7, P8, O1, and O2. EEG signals were recorded with a BrainVision amplifier (Brain Product GmbH, Germany) and sampled at a rate of 1000Hz. Horizontal electrooculogram (HEOG) and vertical electrooculogram (VEOG) were recorded bipolarly from electrodes placed 1cm from the eye. An online reference electrode was placed on the left mastoid and an offline reference electrode was placed on the right mastoid. The ground electrode was placed on the forehead. Impedance was kept below 5 kΩ for all electrodes. EEG recordings took place in an electromagnetically shielded room.

### 2.5 EEG data preprocessing

EEG data were preprocessed and analyzed offline in MATLAB 2021a (The Mathworks, Natick, MA, USA) using the Fieldtrip Toolbox extensions (Oostenveld et al., 2011) as well as custom code. The signal was first segmented into trials from 500 ms pre-stimulus onset (image presentation) to 1500 ms post-stimulus. EEG data were re-referenced to the average of the signal at the left and right mastoids and downsampled to 250 Hz. Ocular movements were removed with Independent Component Analysis (ICA, logistic infomax ICA algorithm; Bell & Sejnowski, 1995); on average, 1.4 ± 0.5 (mean ± SD) eye movement-related components were visually identified and removed per participant. Single trials in which the peak amplitude exceeded 3 SD above/below the mean amplitude were rejected; on average, 91.2 ± 3.4% (mean ± SD) of trials were preserved per participant.

### 2.6 Time-frequency analyses

The preprocessed signal was filtered with a 1-30 Hz bandpass filter. Time-frequency power was computed for each trial by decomposing the signal with a complex Morlet wavelet transformation (frequency-bin size: 1Hz, three cycles per time window; time-bin size: 50 ms). Baseline normalization was performed by log-transforming the power in the epoch of interest (0 −1000 ms post-stimulus) relative to the power in the pre-stimulus interval (500 ∼ 100 ms). The present analysis focuses on power in the theta (4 – 7 Hz) band, based on literature suggesting theta activity plays a role in body processing (see Introduction).

The time window of interest was selected based on previous literature suggesting body-selectivity occurs in the theta band within 250 – 500 ms post-stimulus (Bossi et al., 2020), as well as inspection of the present data, which revealed a peak between 200 – 550 ms post-stimulus for normal compared to scramble conditions (Fig. 2). Based on this observation, the mean theta power during the time window (200 – 550 ms) was extracted at each electrode for all conditions.

**Figure 2.**
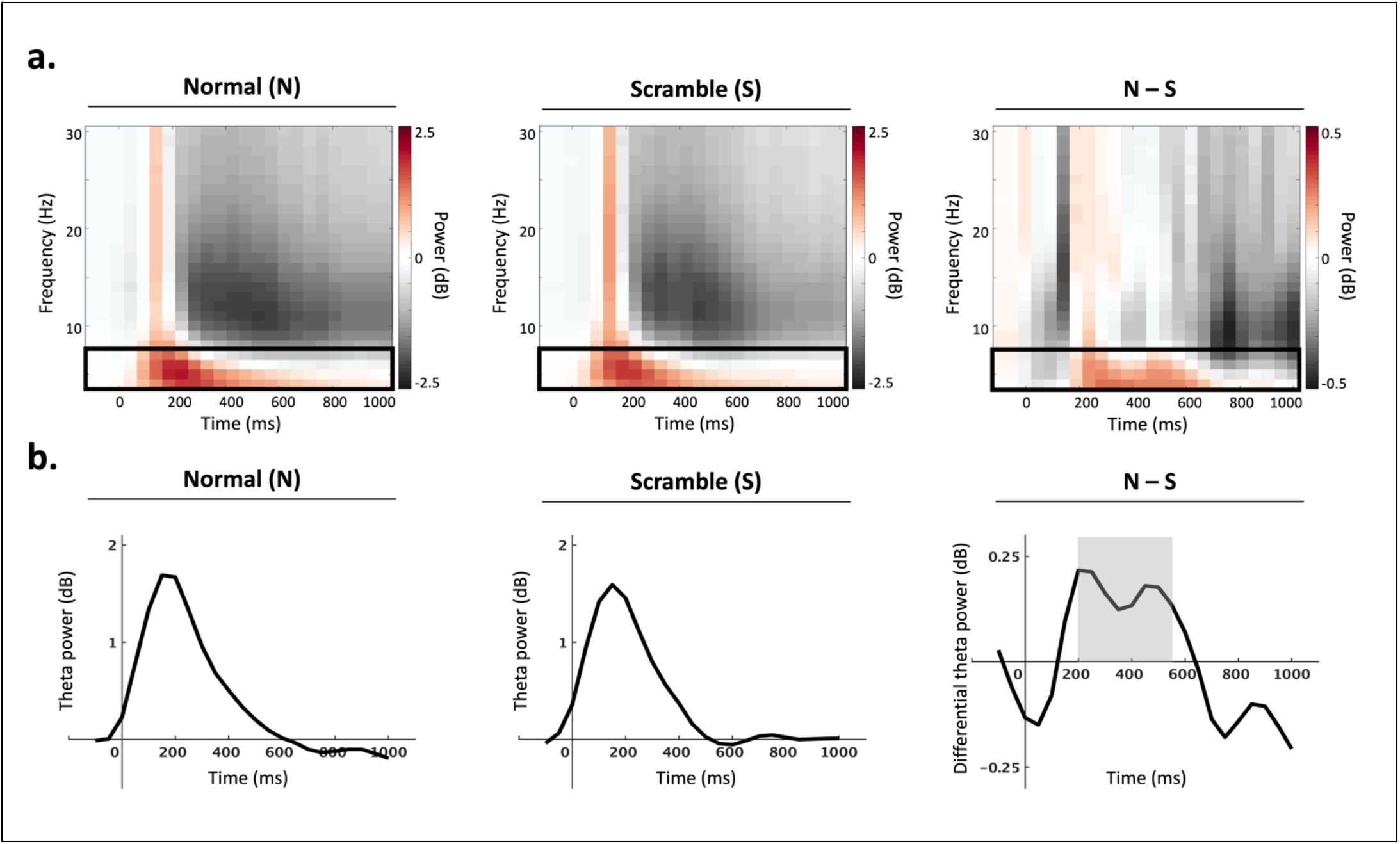
Time window selection. (A) Group-level power spectra computed across all electrodes for all normal (left) and all scramble (middle) conditions. Differential power (normal – scramble) is represented on the right panel. Theta activity (4 – 7 Hz) is indicated with a black box. Power relative to the pre-stimulus baseline is shown in decibels (dB) across time (ms) and frequency (Hz). (B) Time-series of theta power (dB) across conditions. The average theta power computed across all electrodes is shown for all normal (left) and all scramble (middle) conditions.

Differential theta power (normal – scramble) is shown on the right panel, and the time window of interest (200 – 550 ms) is indicated with a grey box.

### 2.7 Cluster-based permutation analyses

To extract regions involved in visual object processing, non-parametric cluster-based permutation analysis was used to select groups of neighboring channels with a significant difference between normal and scramble conditions. With this data-driven method, the mean theta power during the time window of interest (200 – 550 ms) was pooled for all normal (human/monkey * body/face/object) and all scramble (human/monkey * body/face/object) conditions. For each electrode, normal and scramble conditions were compared by means of a t-test (one-sided; normal > scramble). Neighboring electrodes (minimum group size = 2) with t-values exceeding a threshold of p < 0.05 were defined as clusters. Cluster-level test statistics were calculated by summing the t-values within each cluster. To test the statistical significance of the clusters, Monte Carlo permutation tests were run (N = 2,000 permutations) to obtain a null distribution of cluster-level test statistics. Cluster-level test statistics computed from observed data were statistically compared to the reference distribution. Clusters with a probability below a critical alpha level of 0.05 were deemed significant.

Cluster-based permutation analysis of theta power during the time window of interest (200 – 550 ms) revealed a significant difference between normal and scramble conditions in a widespread, bi-lateral cluster, which included 23 electrodes: AFz, FCz, Cz, CPz, Pz, Fp1, Fp2, F3, F4, F7, F8, FC3, FC4, FT7, FT8, C3, C4, CP3, CP4, TP10, P3, P4, and P8 (p = 0.001) (Fig. 3). From this point forward, this group of electrodes is referred to as the scalp region of interest (ROI) and is utilized for further analyses.

**Figure 3.**
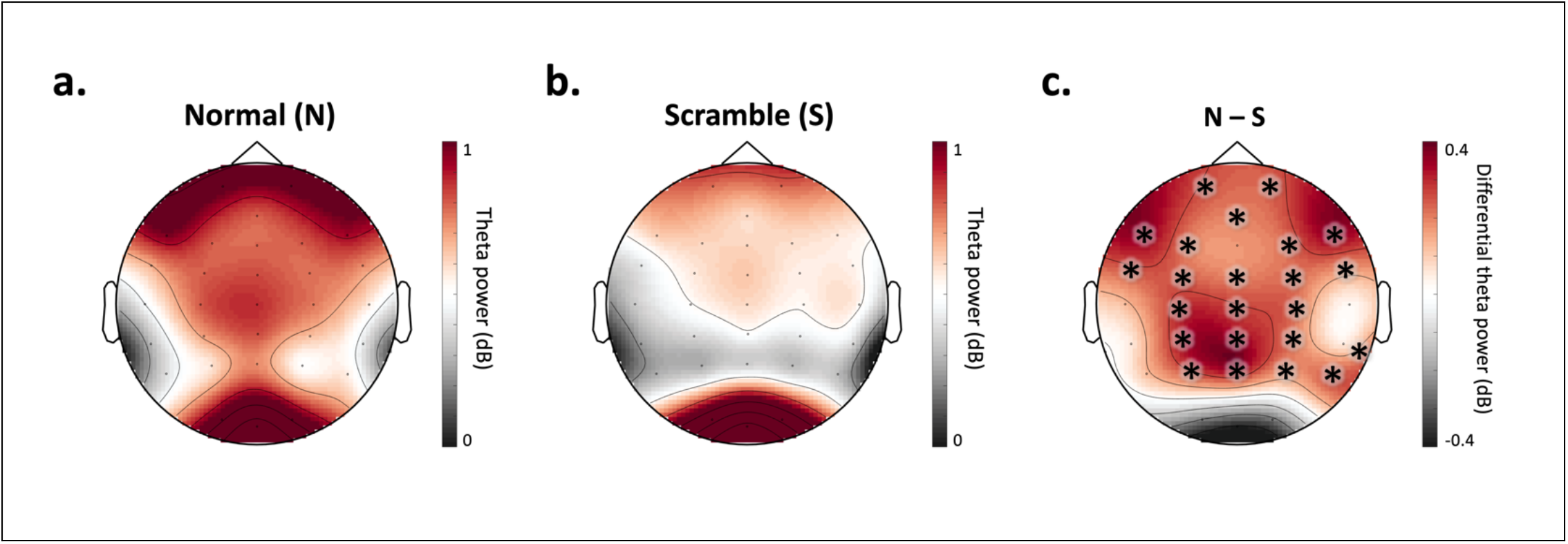
Channel selection. Theta power (4 – 7 Hz) during the time window of interest (200 – 550 ms post-stimulus) for all normal (A) and all scramble (B) conditions. The difference in power (normal – scramble) is represented in (C). Power is shown in decibels (dB). Cluster-based permutation analysis revealed significant differences (p = 0.001) between all normal (A) and all scramble (B) conditions within a cluster of 23 electrodes: AFz, FCz, Cz, CPz, Pz, Fp1, Fp2, F3, F4, F7, F8, FC3, FC4, FT7, FT8, C3, C4, CP3, CP4, TP10, P3, P4, and P8, indicated with asterisks in (C).

### 2.8 Theta power difference

To control for the neural processing of low-level visual features, the difference between normal and scramble conditions was computed for each category. Specifically, the subject-level mean theta activity (200 – 550 ms; ROI) for each scramble condition was subtracted from the respective activity for each normal condition: human body (normal – scramble); monkey body (normal – scramble); human face (normal – scramble); monkey face (normal – scramble); human object (normal – scramble); monkey object (normal – scramble). The resulting differential activity was deemed to represent theta activity related to high-level neural processes and was further analyzed.

### 2.9 Statistical analyses

Statistical analyses were performed using IBM SPSS Statistics 28 (IBM Corp., Armonk, NY, USA). A repeated-measures 2 × 3 ANOVA (Species: human/monkey * Category: body/face/object) was applied to the mean theta power difference (normal – scramble). Statistical differences below p < 0.05 were considered significant. To control for type I errors, a FDR correction was applied to correct for multiple comparisons; only corrected p-values are reported.

## Results

The interaction effect of species*category on differential theta power (normal – scramble) was significant (*F*(2,28) = 4.72, *p* = 0.038, *ηp^2^* = 0.14). The main effect of species (*F*(1,28) = 1.29, *p* = 0.4, *ηp^2^* = 0.04) and the main effect of category (*F* (2,28) = 0.03, *p* = 0.971, *ηp^2^* < 0.001) were not significant. To investigate this interaction effect, three repeated-measures 1 × 2 ANOVAs (Category: bodies × Species: human/monkey; Category: faces × Species: human/monkey; Category: objects × Species: human/monkey) were performed to compare the effect of species on differential theta power (normal – scramble) corresponding to body stimuli, face stimuli and object stimuli, respectively. There was a statistically significant difference in differential theta power between human bodies and monkey bodies (*F*(1,28) = 7.73, *p* = 0.038, *ηp^2^* = 0.22) (Fig. 4-5). Importantly, this species effect was limited to body processing, as no corresponding difference in differential theta power could be found between human faces and monkey faces (*F*(1,28) = 1.74, *p* = 0.395, *ηp^2^* = 0.06), nor between human objects and monkey objects (*F*(1,28) = 0.43, *p* = 0.621, *ηp^2^* = 0.02).

**Figure 4.**
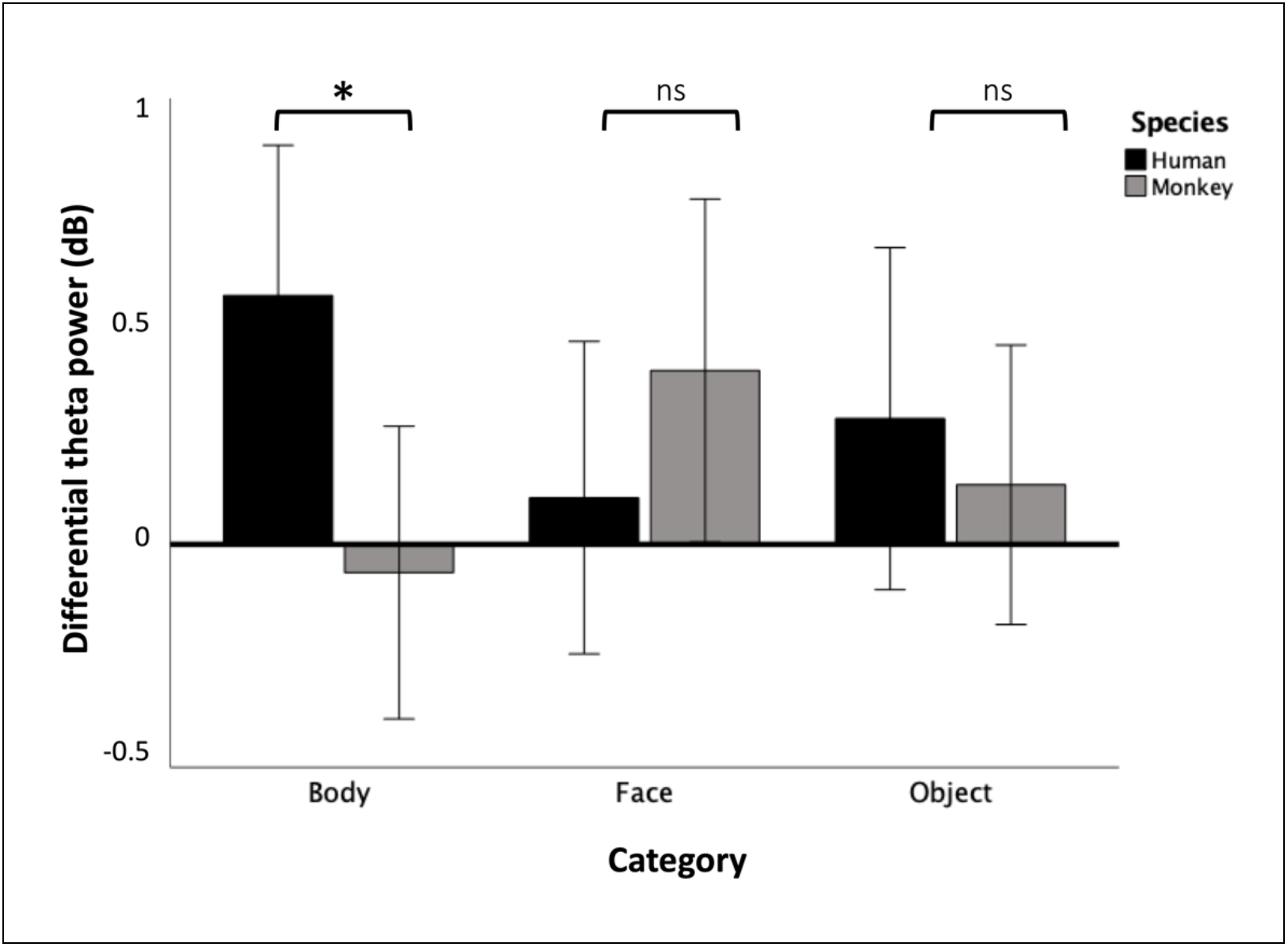
Means of differential theta power (normal – scramble) during the time window of interest (200 – 550 ms post-stimulus), calculated over the ROI for each condition. *: p < 0.05. n.s.: non-significant.

**Figure 5.**
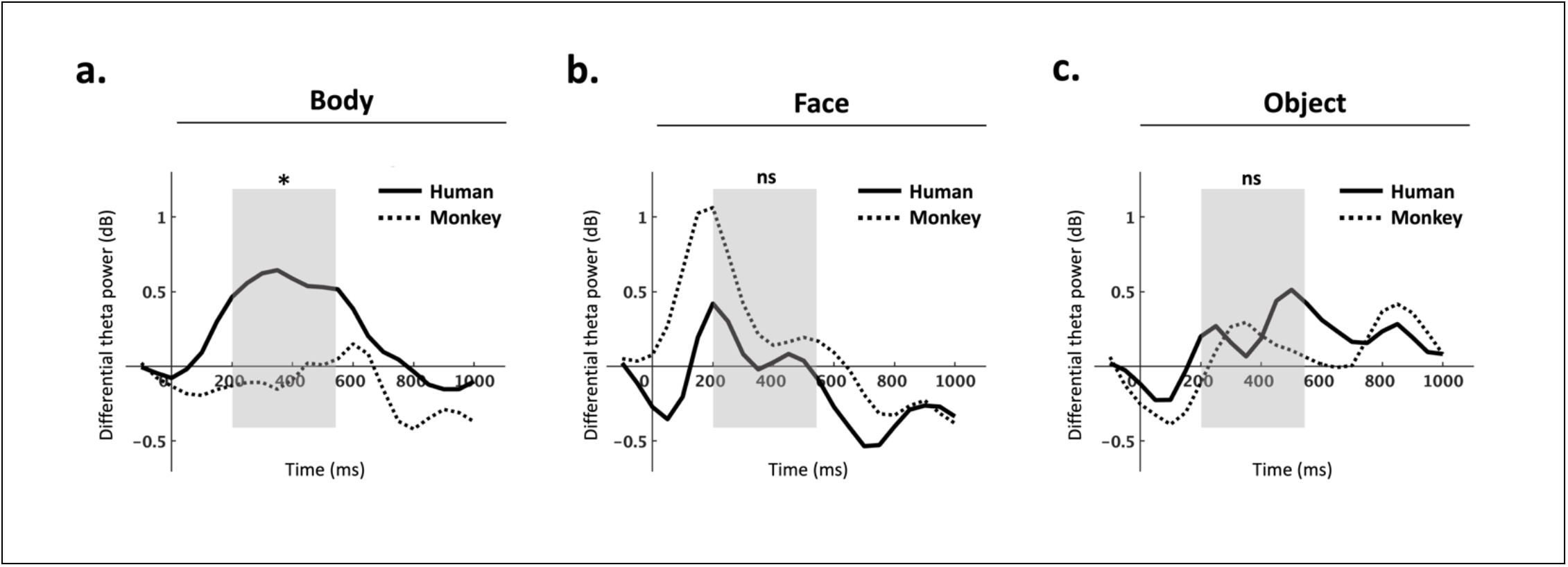
Time-series of differential theta power (normal – scramble) calculated over the ROI, shown separately for body stimuli (A), face stimuli (B), and object stimuli (C). Solid lines represent human stimuli and dashed lines represent monkey stimuli. The time window of interest (200 – 550 ms) is indicated with a grey box. Differential theta power is shown in decibels (dB) and time is shown in milliseconds (ms). Repeated measures ANOVA revealed a significant difference between human body (N-S) and monkey body (N-S) conditions in the time window of interest (p < 0.05) (A), as indicated with an asterisk. This species effect was not significant (ns) among face (B) or object (C) stimuli.

### 3.1 Posthoc analyses and results

Posthoc analyses were run to further characterize the observed effect of species on body processing. First, to explore the spatial distribution of the effect, paired samples t-tests were performed to compare differential theta power between human and monkey body stimuli at each individual channel (N = 33; see Methods). FDR correction was applied to correct for multiple comparisons; only corrected p-values are reported. A significant difference between human body and monkey body in differential theta power was observed at 12 channels within the ROI (AFz, FCz, Cz, CPz, Fp2, F3, FC3, FT7, C3, CP3, P3, and P8) and one channel outside of the ROI (Fz) (p < 0.05; Fig. 6), suggesting that the species effect primarily affected brain regions strongly involved in high-level visual processing. See supplementary materials (Table S3) for results of the individual channel-level paired t-tests.

**Figure 6.**
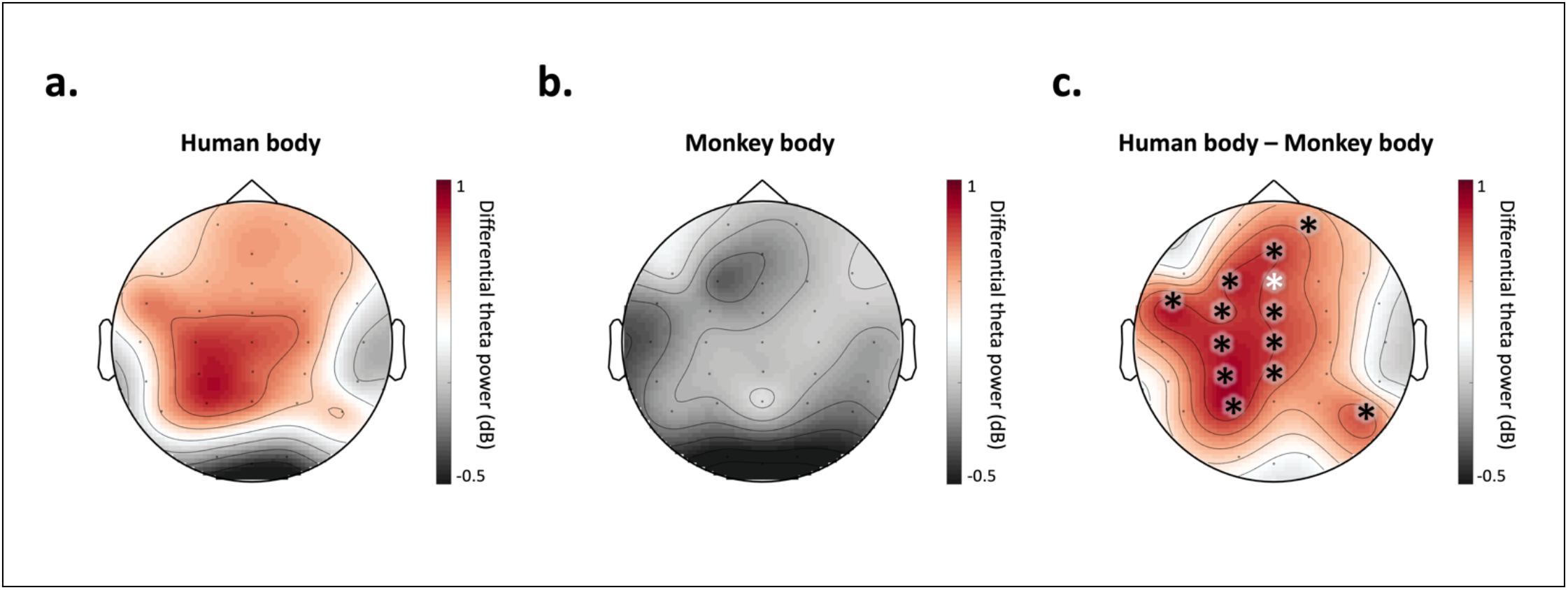
Group-level topography of differential theta power (normal – scramble) during the time window of interest for human body stimuli (A) and monkey body stimuli (B), as well as their difference (C). Asterisks indicate channel locations with a significant difference (p < 0.05) between scramble-controlled human and monkey body conditions, calculated from posthoc paired samples t-tests. Black asterisks represent significant channels that also belong to the ROI (AFz, FCz, Cz, CPz, Fp2, F3, FC3, FT7, C3, CP3, P3, and P8). White asterisks represent channels outside of the ROI (Fz).

Second, to further characterize the temporal profile of the effect of species among body stimuli, temporal cluster-based analysis was performed. Subject-level mean differential theta power in the ROI was computed for human body and monkey body conditions, separately for each time point during the interval 0 to 1000 ms post-stimulus in 50 ms increments (N = 21 time points). These subject-level averages were analyzed with temporal cluster-based analysis, which followed the methodology of the cluster-based analysis used for channel-selection (see Methods), but channels were replaced by time points. Results of the temporal cluster-based analysis of differential theta power in the ROI revealed a significant difference between human body and monkey body at nine consecutive time points between 150 – 550 ms (150, 200, 250, 300, 350, 400, 450, 500 and 550 ms; p = 0.01) (Fig. 7).

**Figure 7.**
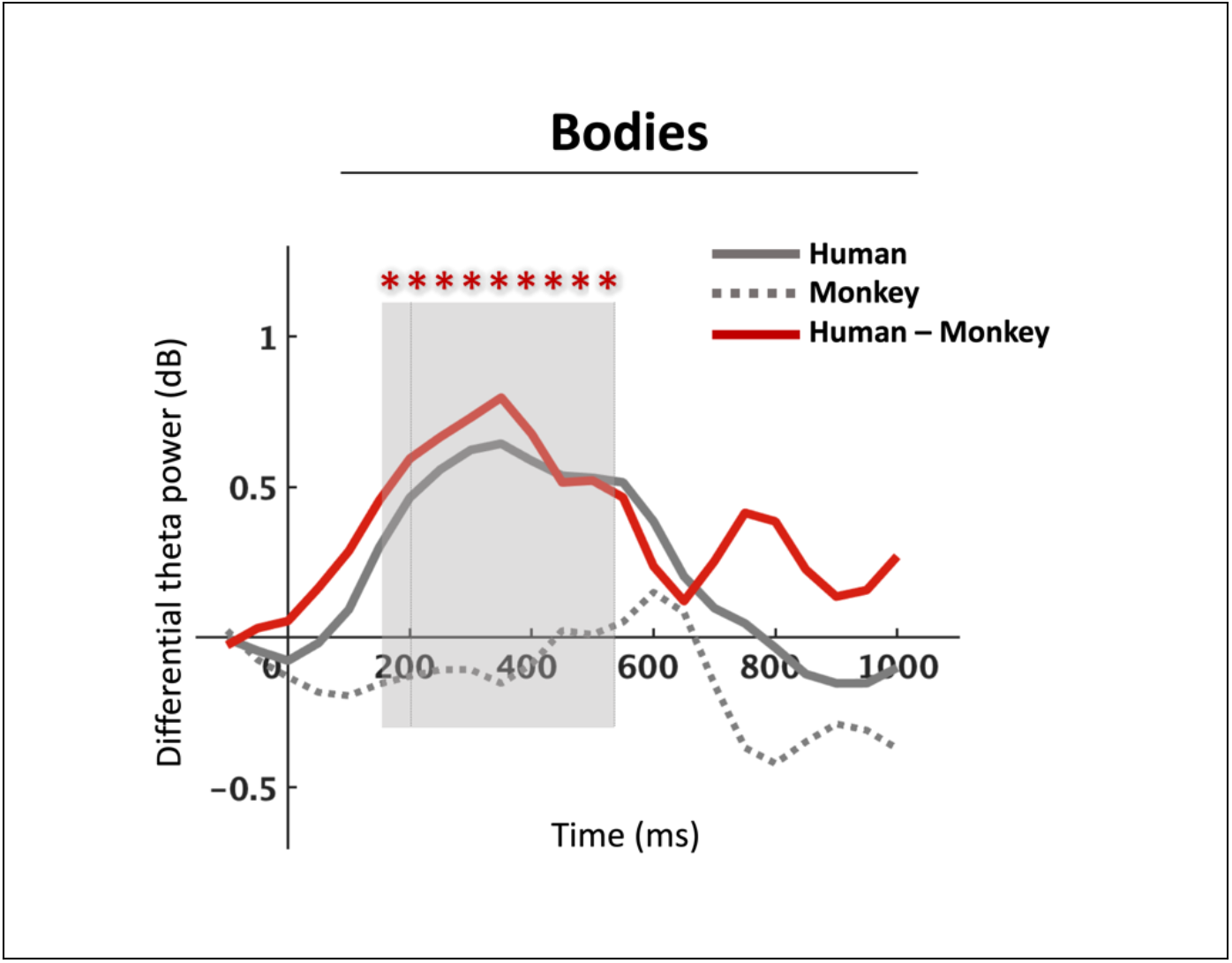
Time-series of differential theta power (normal – scramble) calculated over the ROI, shown separately for human body stimuli (solid gray line), monkey body stimuli (dashed gray line), and their difference (red line). The waveforms corresponding to human body and monkey body stimuli are the same as in Figure 5A. Asterisks represent nine consecutive time points between (150 – 550 ms) with a significant difference (p < 0.05) between scramble-controlled human and monkey body conditions, calculated from posthoc temporal cluster-based analysis. This temporal cluster is highlighted with a grey box. The original time window of interest (200 – 550 ms) is marked with vertical lines.

ERP analyses were performed to further investigate whether the identified oscillatory effect might reflect evoked or induced activity. The same analysis pipeline was applied as for the time-frequency analysis (see Supplementary Analyses). We found no significant difference in ERP amplitude between human bodies and monkey bodies (see Supplementary Results; Fig. S1), mismatching the results based on differential theta power. This indicates that the species effect on body processing was reflected in theta oscillations rather than phase-locked activity.

Finally, to investigate whether the effect was specific to the theta-band, we applied the analysis pipeline to alpha-(8 – 12 Hz) and beta-band (13 – 30 Hz) power (see Supplementary Analyses). There was no significant difference between normal and scramble conditions at any clusters of electrodes during the time window of interest in the alpha-or beta-bands (see Supplementary Results; Fig. S2); no region of interest representing visual object-level processing could be identified.

## 4. Discussion

Our goal was to investigate the time course and functional significance of body processing with a focus on species specificity. We focused on the precise timing and topography of species-specific body processing in the theta-band. Furthermore, given recent fMRI research proposing a large-scale, species-specific cortical network for human body processing (Li et al., 2023), we expected to find a clear enhancement of human (versus monkey) processing. We found a clear effect of species on visual object-level processing that was specific to bodies. More specifically, we found a significant enhancement of the neural representations of human (versus monkey) bodies, and most notably, this species effect was not present among face or object stimuli. This body-specific process affected low-frequency (theta; 4-7Hz) activity likely originating from widespread regions in the cortex during a time window of 150 – 550 ms post-stimulus. Finally, we found this process may reflect induced activity in the theta band, and it did not extend to alpha (8-12 Hz) or beta (13-30 Hz) frequencies. Our findings corroborate previous findings linking oscillatory theta activity to body processing (Bossi et al., 2020; Çelik et al., 2021; Moreau et al., 2020). More importantly, our findings show a specificity of body processing for species, which is consistent with recent fMRI research suggesting body processing is species-specific and topographically widespread beyond EBA (Li et al., 2023; Çelik et al., 2021).

Numerous EEG studies on body processing have focused on the analysis of ERPs, and there is substantial evidence for a body-evoked cortical response at 190 ms (N190) post-stimulus (Peelen & Downing, 2007; Taylor et al., 2010; Thierry et al., 2006; Moreau et al., 2018). On the other hand, oscillatory cortical responses in the context of body processing have been investigated much less, yet the method is powerful in aiding our understanding of cognitive processes reflecting endogenous, non-phase-locked activity, which is attenuated in ERP analyses (Cohen, 2014 & Luck, 2014). Furthermore, modulations of frequency-specific activity have been consistently implicated in cognitive functions (Cavanagh & Frank, 2014; Herweg et al., 2020; Trujillo & Allen, 2006), but only recently have oscillations been investigated in the context of body processing. Recent research has compared theta activation for body versus face processing (Bossi et al., 2020) and self-versus non-self-bodies (Çelik et al., 2021), as well as for body processing amid social interactions (Moreau et al., 2020). Yet, none of these oscillatory studies have investigated species-specific effects, which marks the novelty of the present study.

Our channel-wise exploration of species-specific body processing revealed a bi-lateral cluster, albeit largely on the left-side of the cortex (Fig. 6C). This finding is in line with previous research showing a left-sided effect in the theta band for upright versus inverted bodies (Bossi et al., 2020); this potential left-sided bias is unclear and requires further investigation. In addition, our time point-wise exploration of the precise timing of the species-specific theta effect revealed that the effect emerged from 150 ms and sustained until 550 ms post-stimulus. As our measure of theta activity blended ongoing and phase-locked oscillatory activity, we attempted to separate these two; to this end we analyzed ERPs, a measure of purely phase-locked activity. However, unlike the theta activity-based results, the species-specific effect for bodies in the defined region and time window was not significant in the ERP (see Supplementary Materials; Fig. S1), which may suggest the effect operates on higher-order, top-down processes that are not strictly phase-locked to the visual stimulus (David et al., 2006; Herrmann et al., 2014). Finally, we investigated whether species-specific body processing was reflected in other oscillatory frequency bands, and we did not find any corresponding effect in these oscillatory bands. This further corroborates previous research suggesting oscillatory theta activity plays a relevant role in body processing (Bossi et al., 2020; Çelik et al., 2021; Moreau et al., 2020). Nevertheless, it’s possible oscillatory activity in other frequency bands may also play a role in body processing, and an interesting future direction can investigate those effects in other time-windows.

So far, species-specificity is not fully understood in the nonhuman primate brain. There is consistent evidence for body-selective patches in the macaque temporal cortex (for a review, see Vogels, 2022). In addition, single-unit recordings directly from body-selective patches in the macaque STS revealed differences between bodies and non-bodies, as well as between humans and monkeys, indicating effects at multiple processing levels (Kumar & Vogels, 2019). A follow-up to the present study can address the generalizability of our findings to nonhuman primate observers of primate bodies. More specifically, we would expect to find that in the nonhuman primate cortex, theta activity is enhanced in response to images of monkey versus human bodies. An additional future direction can integrate the findings of human and monkey studies to create a comprehensive model of body processing in the brain. Recently, neural network models (Kumar et al., 2023) and theoretical frameworks (de Gelder & Poyo Solanas, 2021) for body processing have been proposed, but we do not have a complete understanding of the neural representations of bodies (Vogels, 2022).

A central question concerns the functional significance of theta oscillations associated with species-specific body processing. Recent reports of theta oscillations offer some interesting and suggestive indications. Studies involving simple conflict paradigms have long suggested theta activity is a mechanism for cognitive control (for a review, see Cavanagh & Frank, 2014). More recently, theta activity was measured in response to approach-avoidance conflicts for the first time, and findings showed a direct relationship between midfrontal theta activation and approach-avoidance conflicts (Lange et al., 2022). A different but potentially highly relevant role of theta oscillations concerns perception-movement initiation at early stages. For example, oscillations in the theta-band may play an important role in combining in a common temporal reference frame visual perception and motor intention (Tomassini et al., 2017). Furthermore, studies on body perception have systematically shown that observing whole body actions is associated with activity in premotor and motor areas (de Gelder et al., 2010; Grèzes et al., 2007; Goldberg et al., 2014; Pichon et al., 2009). The theta effects observed in the present study may be linked to visual body perception in combination with processes related to movement intention. This pattern may have been driven by the inclusion of threatening stimuli, reflecting well-established processes seen in the theta band and related to cognitive control (for a review, see Cavanagh & Frank, 2014). The images used in the present design were selected to have a wide range of body expressions, including neutral expressions as well as emotional expressions depicting defensive actions (fear) and aggressive actions (fear), among others. This does not reduce the importance of the species-specific effect, as the monkey stimulus set equally included neutral and emotionally expressive actions but did not show a similar theta response. Taken together, the observed theta band activity provides clear suggestions for the underlying functional significance of species-specificity.

Another key feature of bodies is dynamics. In daily life, people who interact are not stationary but rather they are, to some degree, always moving. Emerging research using dynamic body stimuli has shown body-and motion-selective processes may be integrated (Raman et al., 2023; Kumar et al., 2023). While the present study used static images, future research should implement dynamic videos to understand the full extent of oscillatory representations of social interactions beyond static object recognition.

## 5. Data availability

The data that support the findings of this study are available on request from the corresponding author (B.d.G), pending approval from the researcher’s local ethics committee and a formal data sharing agreement.

## 6. Author contributions

Jane Chesley (Conceptualization, Data curation, Formal analysis, Investigation, Methodology, Visualization, Writing – original draft, Writing – review & editing, Project administration), Lars Riecke (Conceptualization, Methodology, Writing – review & editing, Supervision), Juanzhi Lu (Formal analysis, resources), Rufin Vogels (Writing – review & editing), Beatrice de Gelder (Conceptualization, Methodology, Writing – review & editing, Supervision, Funding acquisition).

## 9. Supplementary Materials for

### 9.1 Supplementary Analyses

#### 9.1.1 Event-related potential analyses

ERP analyses were performed to further investigate whether the oscillatory effect might reflect evoked or induced activity. Here, the preprocessed EEG signal was baseline-corrected by subtracting the average amplitude during the interval (– 200 ∼ 0 ms) pre-stimulus, and a 50 Hz notch filter was applied. For each condition, the grand-averaged ERP was calculated over the cluster (n = 13; AFz, Fz, FCz, Cz, CPz, Fp2, F3, FC3, FT7, C3, CP3, P3, and P8) identified in the posthoc time-frequency analyses as having a significant species-effect among body stimuli (Fig. 6C). To control for the neural processing of low-level visual features, the amplitude difference (normal – scramble) was calculated for each condition. The mean amplitude difference within the cluster and during the time window (150 – 550 ms) identified in the posthoc time-frequency analyses (Fig. 7) was statistically analyzed with the same repeated measures ANOVAs as for the time-frequency analysis; see Statistical Analyses.

#### 9.1.2 Time-frequency analyses: Alpha-and beta-band activity

Finally, to investigate whether the effect was specific to the theta-band, we applied the analysis pipeline to alpha-(8-12 Hz) and beta-band (13-30 Hz) power. Alpha-and beta-band power during the time window of interest was extracted from the preprocessed, time-frequency transformed signal (see above). Then, to localize object-level processing channels, cluster-based permutation analysis was applied to compare all normal and all scramble conditions.

### 9.2 Supplementary Results

#### 9.2.1 Event-related potential results

In line with the results based on differential theta power, the interaction effect of species*category was significant (*F*(2,28) = 16.28, *p* = 0.003, *ηp^2^* = 0.37). The main effect of species (*F*(1,28) = 8.57, *p* = 0.014, *ηp^2^* = 0.23) was significant and the main effect of category (*F*(2,28) = 0.61, *p* = 0.659, *ηp^2^* = 0.02) was not significant. While there was a statistically significant difference in amplitude between human faces and monkey faces (*F*(1,28) = 27.37, *p* = 0.003, *ηp^2^* = 0.49), there was no significant difference in amplitude between human bodies and monkey bodies (*F*(1,28) = 0.004, *p* = 0.948, *ηp^2^* = 0), nor between human objects and monkey objects (*F*(1,28) = 1.96, *p* = 0.26, *ηp^2^* = 0.07), mismatching the results based on differential theta power (Fig. S1). This indicates that the species effect on body processing was reflected in theta oscillations rather than stimulus phase-locked activity.

#### 9.2.2 Time-frequency results: Alpha-and beta-band activity

There was no significant difference between normal and scramble conditions at any clusters of electrodes during the time window of interest in the alpha-or beta-bands (Fig. S2).

### 9.3 Supplementary Figures and Tables

**Figure S1.**
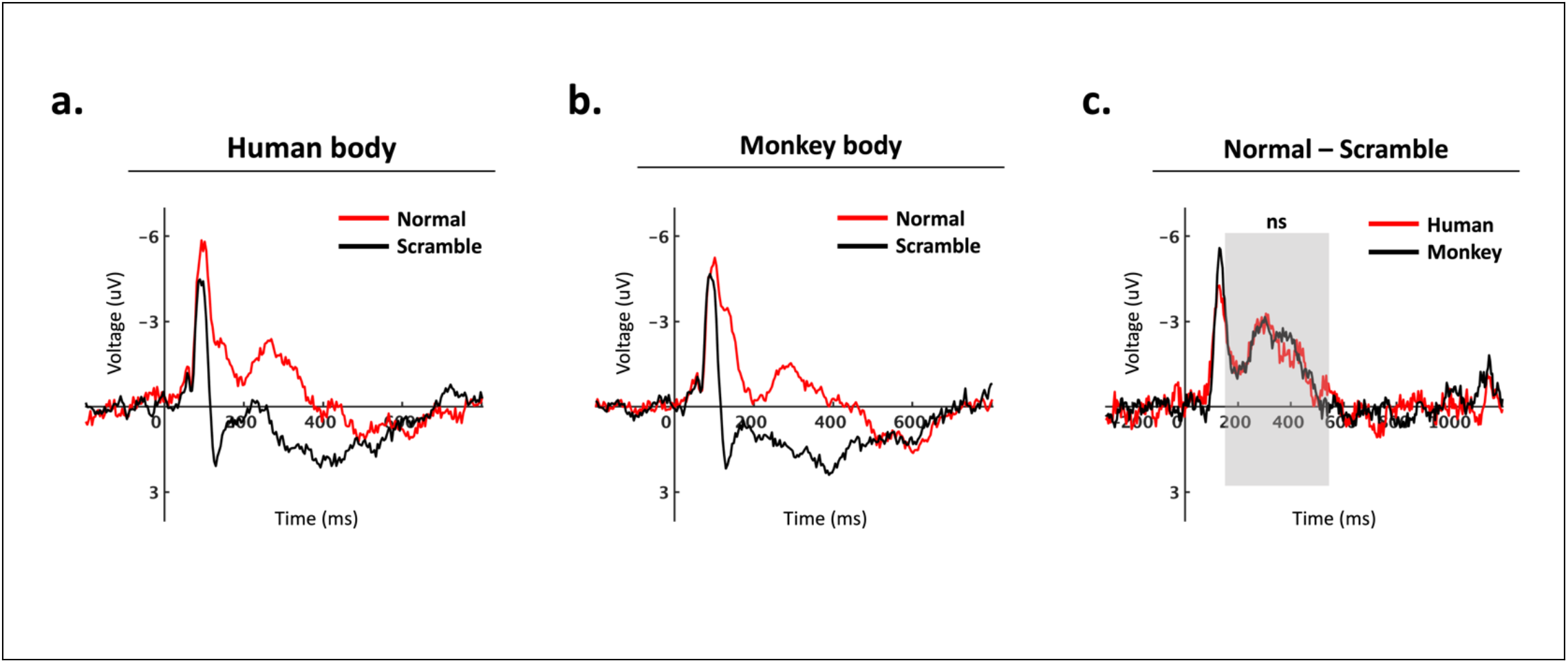
Grand-averaged ERP waveforms per condition (A – B), calculated by averaging the data at electrodes AFz, Fz, FCz, Cz, CPz, Fp2, F3, FC3, FT7, C3, CP3, P3, and P8. (C) Difference waveforms (normal – scramble) shown separately for human and monkey body stimuli. The grey box highlights the time window (150 – 550 ms) used for statistical analyses. There was no significant difference between scramble-controlled human and monkey body stimuli.

**Figure S2.**
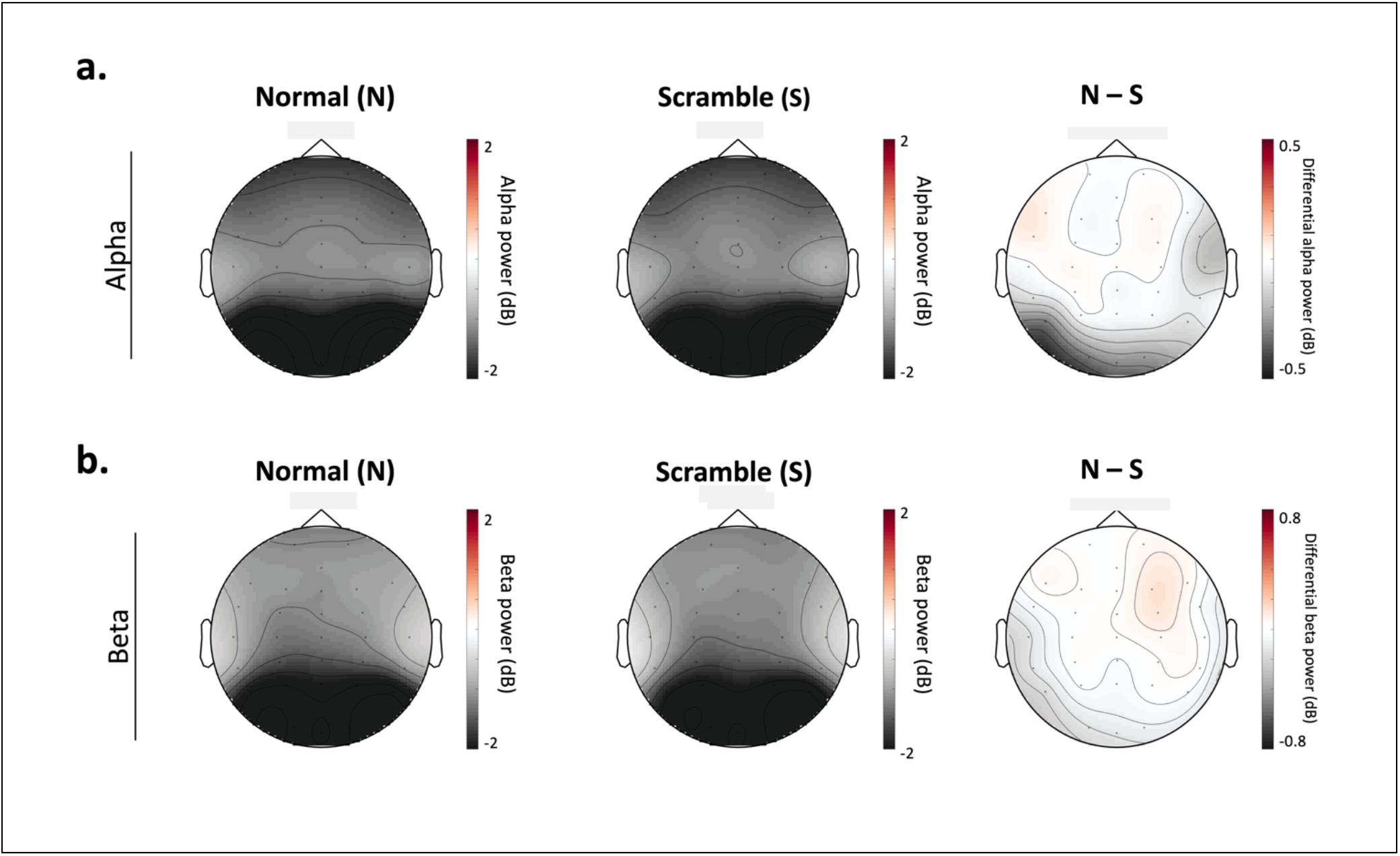
(A) Alpha power (8 – 12 Hz) and (B) beta power (13 – 30 Hz) during the time window of interest (200 – 550 ms post-stimulus) for all normal (left) and all scramble (middle) conditions. The difference in power (normal – scramble) is represented on the right. Cluster-based permutation analysis revealed no significant difference between all normal and all scramble conditions within any clusters of electrodes in alpha-or beta-band frequencies.

**Table S3.** Individual channel-level results of paired t-tests comparing differential theta power (normal – scramble) between human body and monkey body stimuli within the time window of interest (200 – 550 ms). Only significant effects (p < 0.05) with FDR correction are reported.

| Channel | Human body (Normal – Scramble) | Monkey body (Normal – Scramble) | T-test |
| --- | --- | --- | --- |
| AFz | human (M = 0.58, SD = 1.35) | monkey (M = -0.21, SD = 1.17) | t(28) = 2.609, p = 0.040 |
| Fz | human (M = 0.54, SD = 1.29) | monkey (M = -0.19, SD = 1.11) | t(28) = 2.293, p = 0.041 |
| FCz | human (M = 0.63, SD = 1.25) | monkey (M = -0.11, SD = 1.09) | t(28) = 2.267, p = 0.041 |
| Cz | human (M = 0.75, SD = 1.20) | monkey (M = -0.03, SD = 1.14) | t(28) = 2.256, p = 0.041 |
| CPz | human (M = 0.70, SD = 1.18) | monkey (M = -0.03, SD = 1.03) | t(28) = 2.299, p = 0.041 |
| Fp2 | human (M = 0.53, SD = 1.40) | monkey (M = -0.06, SD = 1.35) | t(28) = 2.276, p = 0.041 |
| F3 | human (M = 0.52, SD = 1.29) | monkey (M = -0.29, SD = 1.14) | t(28) = 2.761, p = 0.040 |
| FC3 | human (M = 0.61, SD = 1.20) | monkey (M = -0.13, SD = 1.13) | t(28) = 2.429, p = 0.041 |
| FT7 | human (M = 0.63, SD = 1.19) | monkey (M = -0.20, SD = 1.24) | t(28) = 3.072, p = 0.040 |
| C3 | human (M = 0.79, SD = 1.09) | monkey (M = -0.05, SD = 1.21) | t(28) = 2.718, p = 0.040 |
| CP3 | human (M = 0.82, SD = 1.17) | monkey (M = -0.03, SD = 1.20) | t(28) = 2.687, p = 0.040 |
| P3 | human (M = 0.81, SD = 1.23) | monkey (M = -0.09, SD = 1.21) | t(28) = 2.865, p = 0.040 |
| P8 | human (M = 0.44, SD = 1.03) | monkey (M = -0.25, SD = 1.07) | t(28) = 2.469, p = 0.041 |

## References

1. Bell, A. J., & Sejnowski, T. J. (1995). An information-maximization approach to blind separation and blind deconvolution. Neural Computation, 7(6), 1129–1159. 10.1162/neco.1995.7.6.1129.

2. Brainard, D.H. (1997). The Psychophysics Toolbox. Spatial Vision, 10(4):433–6.

3. Bognár, A., Raman, R., Taubert, N., Zafirova, Y., Li, B., Giese, M., De Gelder, B., & Vogels, R. (2023). The contribution of dynamics to macaque body and face patch responses. NeuroImage, 269, 119907. 10.1016/j.neuroimage.2023.119907.

4. Bossi, F., Premoli, I., Pizzamiglio, S., Balaban, S., Ricciardelli, P., & Rivolta, D. (2020). Theta-and Gamma-Band Activity Discriminates Face, Body and Object Perception. Frontiers in Human Neuroscience, 14. 10.3389/fnhum.2020.00074.

5. Candidi, M., Stienen, B. M., Aglioti, S. M., & de Gelder, B. (2015). Virtual lesion of right posterior superior temporal sulcus modulates conscious visual perception of fearful expressions in faces and bodies. Cortex, 65, 184–194. 10.1016/j.cortex.2015.01.012.

6. Cavanagh, J. F., & Frank, M. J. (2014). Frontal theta as a mechanism for cognitive control. Trends in Cognitive Sciences, 18(8), 414–421. 10.1016/j.tics.2014.04.012.

7. Çelik, S., Doğan, R. B., Parlatan, C. S., et al. (2021). Distinct brain oscillatory responses for the perception and identification of one’s own body from other’s body. Cognitive Neurodynamics, 15(6), 609–620. 10.1007/s11571-020-09660-z.

8. Cohen, M. X. (2014). Analyzing Neural Time Series Data: Theory and Practice. *The MIT Press*. 10.7551/mitpress/9609.001.0001.

9. David, O., Kilner, J., & Friston, K. (2006). Mechanisms of evoked and induced responses in MEG/EEG. NeuroImage, 31, 1580–1591. 10.1016/j.neuroimage.2006.02.034.

10. de Gelder, B., & Poyo Solanas, M. (2021). A computational neuroethology perspective on body and expression perception. Trends in Cognitive Sciences, 25(9), 744–756. 10.1016/j.tics.2021.05.010.

11. de Gelder, B., Van den Stock, J., Meeren, H. K. M., Sinke, C. B. A., Kret, M. E., & Tamietto, M. (2010). Standing up for the body: Recent progress in uncovering the networks involved in the perception of bodies and bodily expressions. Neuroscience & Biobehavioral Reviews, 34(4), 513–527. 10.1016/j.neubiorev.2009.10.008.

12. Downing, P. E., Jiang, Y., Shuman, M., & Kanwisher, N. (2001). A Cortical Area Selective for Visual Processing of the Human Body. Science, 293(5539), 2470–2473. http://www.jstor.org/stable/3084903.

13. Freiwald, W., Duchaine, B., & Yovel, G. (2016). Face Processing Systems: From Neurons to Real-World Social Perception. Annual Review of Neuroscience, 39(1), 325–346. 10.1146/annurev-neuro-070815-013934.

14. Fries, P. (2009). Neuronal gamma-band synchronization as a fundamental process in cortical computation. Annual Review of Neuroscience, 32, 209–224. 10.1146/annurev.neuro.051508.135603.

15. Fries, P. (2015). Rhythms for Cognition: Communication through Coherence. Neuron, 88(1), 220–235. 10.1016/j.neuron.2015.09.034.

16. Goldberg, H., Preminger, S., & Malach, R. (2014). The emotion-action link? Naturalistic emotional stimuli preferentially activate the human dorsal visual stream. Neuroimage, 84, 254–264. 10.1016/j.neuroimage.2013.08.032.

17. Grèzes, J., Pichon, S., & de Gelder, B. (2007). Perceiving fear in dynamic body expressions. Neuroimage, 35(2), 959–967. 10.1016/j.neuroimage.2006.11.030.

18. Groen, I. I. A., Silson, E. H., & Baker, C. I. (2017). Contributions of low-and high-level properties to neural processing of visual scenes in the human brain. Royal Society. 10.1098/rstb.2016.0102.

19. Herrmann, C. S., Rach, S., Vosskuhl, J., et al. (2014). Time–Frequency Analysis of Event-Related Potentials: A Brief Tutorial. Brain Topography, 27, 438–450. 10.1007/s10548-013-0327-5.

20. Herweg, N. A., Solomon, E. A., & Kahana, M. J. (2020). Theta Oscillations in Human Memory. Trends in Cognitive Sciences, 24(3), 208–227. 10.1016/j.tics.2019.12.006.

21. Kandel, E. R., Schwartz, J. H., Jessell, T. M., Siegelbaum, S. A., Hudspeth, A. J., & Mack, S. (Eds.) (2014). High-level visual processing: cognitive influences. *In Principles of Neural Science, Fifth Edition. McGraw Hill*. https://neurology.mhmedical.com/content.aspx?bookid=1049&sectionid=59138656.

22. Kleiner, M., Brainard, D., & Pelli, D. (2007). What’s new in Psychtoolbox-3? *Perception, 36*, ECVP Abstract Supplement.

23. Kret, M. E., Pichon, S., Grèzes, J., & de Gelder, B. (2011). Similarities and differences in perceiving threat from dynamic faces and bodies: An fMRI study. Neuroimage, 54(2), 1755–1762. 10.1016/j.neuroimage.2010.08.012.

24. Koch, C., Ullman, S. (1987). Shifts in Selective Visual Attention: Towards the Underlying Neural Circuitry. In: Vaina, L.M. (eds) Matters of Intelligence. Synthese Library, vol 188. Springer, Dordrecht. 10.1007/978-94-009-3833-5_5.

25. Kumar, S., & Vogels, R. (2019). Body Patches in Inferior Temporal Cortex Encode Categories with Different Temporal Dynamics. Journal of Cognitive Neuroscience, 31(11), 1699–1709. 10.1162/jocn_a_01444.

26. Kumar, P. et al. (2023). Neurodynamical Model of the Visual Recognition of Dynamic Bodily Actions from Silhouettes. In: Iliadis, L., Papaleonidas, A., Angelov, P., Jayne, C. (eds) Artificial Neural Networks and Machine Learning – ICANN 2023. ICANN 2023. Lecture Notes in Computer Science, vol 14255. Springer, Cham. 10.1007/978-3-031-44210-0_43.

27. Lange, L., Rommerskirchen, L., & Osinsky, R. (2022). Midfrontal Theta Activity Is Sensitive to Approach–Avoidance Conflict. Journal of Neuroscience, 42(41), 7799–7808. 10.1523/JNEUROSCI.2499-21.2022.

28. Li, B., Poyo Solanas, M., Marrazzo, G., Raman, R., Taubert, N., Giese, M., Vogels, R., & de Gelder, B. (2023). A large-scale brain network of species-specific dynamic human body perception. Progress in Neurobiology, 221, 102398. 10.1016/j.pneurobio.2022.102398.

29. Luck, S. J. (2014). *An Introduction to the Event-Related Potential Technique, Second Edition*. The MIT Press. ISBN: 9780262525855.

30. Moreau, Q., Candidi, M., Era, V., Tieri, G., & Aglioti, S. M. (2020). Midline frontal and occipito-temporal activity during error monitoring in dyadic motor interactions. Cortex, 127, 131–149. 10.1016/j.cortex.2020.01.020.

31. Moreau, Q., Pavone, E. F., Aglioti, S. M., & Candidi, M. (2018). Theta synchronization over occipito-temporal cortices during visual perception of body parts. European Journal of Neuroscience, 48(9), 2826–2835. 10.1111/ejn.13782.

32. Oostenveld, R., Fries, P., Maris, E., & Schoffelen, J. M. (2011). FieldTrip: Open Source Software for Advanced Analysis of MEG, EEG, and Invasive Electrophysiological Data.

33. Peelen, M. V., & Downing, P. E. (2005). Selectivity for the Human Body in the Fusiform Gyrus. Journal of Neurophysiology, 93(1), 603–608. 10.1152/jn.00513.2004.

34. Peelen, M. V., & Downing, P. E. (2007). The neural basis of visual body perception. Nature Reviews Neuroscience, 8(8), 636–648. 10.1038/nrn2195.

35. Pelli, D. G. (1997). The VideoToolbox software for visual psychophysics: Transforming numbers into movies. Spatial Vision, 10(4), 437–442. 10.1163/156856897X00366

36. Pichon, S., de Gelder, B., & Grèzes, J. (2009). Two different faces of threat: comparing the neural systems for recognizing fear and anger in dynamic body expressions. NeuroImage, 47(4), 1873–1883. 10.1016/j.neuroimage.2009.03.084.

37. Pourtois, G., Peelen, M. V., Spinelli, L., Seeck, M., & Vuilleumier, P. (2007). Direct intracranial recording of body-selective responses in human extrastriate visual cortex. Neuropsychologia, 45(11), 2621–2625. 10.1016/j.neuropsychologia.2007.04.005.

38. Powell, L. J., Kosakowski, H. L., & Saxe, R. (2018). Social Origins of Cortical Face Areas. Trends in Cognitive Sciences, 22(9), 752–763. 10.1016/j.tics.2018.06.009.

39. Poyo Solanas, M., Vaessen, M., & de Gelder, B. (2020). Computation-Based Feature Representation of Body Expressions in the Human Brain. Cerebral Cortex, 30(12), 6376– 6390. 10.1093/cercor/bhaa196.

40. Raman, R., Bognar, A., Ghamkari Nejad, G., Taubert, N., Giese, M., & Vogels, R. (2023). Bodies in Motion: Unraveling the Distinct Roles of Motion and Shape to Dynamic Body Responses in Temporal Cortex. *Cell reports*. (in press).

41. Schwiedrzik, C. M., Zarco, W., Everling, S., & Freiwald, W. A. (2015). Face Patch Resting State Networks Link Face Processing to Social Cognition. PLOS Biology. 10.1371/journal.pbio.1002245.

42. Stekelenburg, J. J., & de Gelder, B. (2004). The neural correlates of perceiving human bodies: an ERP study on the body-inversion effect. Neuroreport, 15, 777–780. doi:10.1097/01.wnr.0000119730.93564.e8.

43. Swann, P., Pichon, S., de Gelder, B., & Grèzes, J. (2012). Threat prompts defensive brain responses independently of attentional control. Cerebral Cortex, 22(2), 274–285. 10.1093/cercor/bhr060.

44. Taylor, J. C., Roberts, M. V., Downing, P. E., & Thierry, G. (2010). Functional characterisation of the extrastriate body area based on the N1 ERP component. Brain and Cognition, 73(3), 153–159. 10.1016/j.bandc.2010.04.001.

45. Thierry, G., Pegna, A. J., Dodds, C., Roberts, M., Basan, S., & Downing, P. (2006). An event-related potential component sensitive to images of the human body. NeuroImage, 32(2), 871–879. 10.1016/j.neuroimage.2006.03.060.

46. Tomassini, A., Ambrogioni, L., Medendorp, W. P., & Maris, E. (2017). Theta oscillations locked to intended actions rhythmically modulate perception. eLife, 6, e25618. 10.7554/eLife.25618.

47. Trujillo, L. T., & Allen, J. J. B. (2007). Theta EEG dynamics of the error-related negativity. Clinical Neurophysiology, 118(3), 645–668. 10.1016/j.clinph.2006.11.009.

48. van Heijnsbergen, C. C., Meeren, H. K., Grèzes, J., & de Gelder, B. (2007). Rapid detection of fear in body expressions, an ERP study. Brain Research, 1186, 233–241. 10.1016/j.brainres.2007.09.093.

49. Veale, R., Hafed, Z. M., & Yoshida, M. (2017). How is visual salience computed in the brain? Insights from behaviour, neurobiology and modelling. The Royal Society, 372(1714), 20160113. 10.1098/rstb.2016.0113.

50. Vogels, R. (2022). More Than the Face: Representations of Bodies in the Inferior Temporal Cortex. Annual Review of Vision Science, 8, 383–405. 10.1146/annurev-vision-100720-113429.

51. Wang, X. J. (2010). Neurophysiological and Computational Principles of Cortical Rhythms in Cognition. Physiological Reviews, 90(3), 1195–1268. 10.1152/physrev.00035.2008.

52. Zhu, Q., Nelissen, K., Van den Stock, J., De Winter, F. L., Pauwels, K., de Gelder, B., Vanduffel, W., & Vandenbulcke, M. (2013). Dissimilar processing of emotional facial expressions in human and monkey temporal cortex. NeuroImage, 66, 402–411. 10.1016/j.neuroimage.2012.10.083.

